# Technical report on best practices for hybrid and long read *de novo* assembly of bacterial genomes utilizing Illumina and Oxford Nanopore Technologies reads

**DOI:** 10.1101/2022.10.25.513682

**Authors:** Simon T. Hackl, Theresa A. Harbig, Kay Nieselt

**Affiliations:** Institute for Bioinformatics and Medical Informatics, University of Tübingen, 72076, Tübingen, Germany

## Abstract

The emergence of commercial long read sequencing technologies in the 2010s and the concomitant development of new bioinformatics tools bears the potential of *de novo* genome assemblies of unprecedented contiguity and quality. However, until today these novel technologies suffer from high rates of sequencing errors. These may be overcome by using long and short reads in combination, in so called hybrid approaches, or by increasing the throughput and thereby the coverage of sequencing runs. In particular the latter will thereby increase the cost of the assembly inevitably. Herein, to-date long read and hybrid assemblers were tested on real whole genome sequencing Illumina and Oxford Nanopore Technologies read data sets and sub samples of these in order to elaborate a best practice for *de novo* assembly. The findings suggest that although long reads alone can be used to reconstruct complete and contiguous genomes, in particular the single-nucleotide and indel error rate remains high compared to hybrid approaches and that this can impact downstream applications such as variation discovery and gene prediction negatively.

## Introduction

Over the past four decades DNA sequencing has experienced a history rich in technological breakthroughs [1]. Yet, modern devices are unable to sequence a full-length genome without errors in a single stretch and thus one remaining fundamental problem of bioinformatics is *de novo* genome assembly [2]. This problem may informally be described as the reconstruction of as much of a genome sequence as possible given a collection of genomic fragments, so called reads, without a reference genome [2].

Bioinformatics tools used to solve this problem are referred to as assemblers and commonly produce a set of contigs, contiguous stretches of a genome, from overlapping reads [2]. In an optimal scenario one error free contig per chromosome or plasmid of the organism of interest would have to be derived.

State of the art technologies used to generate huge amounts of read data at an appropriate cost may be classified as amplification-based- or *second generation sequencing* (SGS) and single-molecule- or *third generation sequencing* (TGS) [3]. A comprehensive overview of the single technologies in each class is given for example by Schendure *et al*. [1]. Herein, the focus lies on Illuminas SGS and *Oxford Nanopore Technologies* (ONT) TGS platforms.

A noteable hallmark in the development of SGS is the acquisition of market dominance by Illumina around 2012 and an associated saturation of the technology in terms of accuracy, data throughput and cost [1]. Segerman reports that in 2020 82 % of all bacterial genomes in the RefSeq database, a database containing high-quality bacterial assemblies maintained by the *National Center for Biotechnology Information* (NCBI), were produced using the Illuminas SGS technology [4].

ONT released their first TGS devices in 2014 [1]. In 2020, according to Segermans evaluation of the RefSeq database, ONT sequencing constitutes only 3 % of all stored sequences, however its usage rate is gradually increasing [4]. Its main key feature is the generation of long reads spanning thousands and millions of base pairs, yet they yield comparatively high error rates [5].

A trending development is the combination of both technologies for *de novo* assembly. These hybrid assembly approaches commonly pair the information about the global genomic structure, being contained in long reads, with the high accuracy of short reads. Approximately 90 % of all sequences completed during 2019 are based on a combination of Illumina and TGS technologies of which 75 % are accounted to the combination of Illumina and ONT sequencing [4]. Especially bacterial genome assemblies are fostered by the rise of TGS may be accounted to the simpler structure of such regarding size, repetitive patterns and ploidity [6].

Still, the generation of high coverage whole genome sequencing data with ONT devices is cost-intensive [5]. Obliviously, one mostly seeks for a study design that ensures robust results for the lowest cost.

This development of new sequencing technologies goes hand-in-hand with the emergence of a multitude of new bioinformatics tools, not only used for *de novo* assembly per se, but also concerned with the correction of errors in reads and contigs [7].

These changes in the landscape of genome sequencing inevitably raise the questions of what computational approach and which sequencing technology yield the best result and how much data is needed in order to run valid long read only as well as hybrid assemblies.

In order to address these questions this project aims at (i) investigating performance differences of assemblers between the class of hybrid and long read assemblers, but also within these classes, using real prokaryotic read data sets, (ii) elaborating the effect of decreasing long read coverage on the assembly quality and (iii) inspecting the possible application of long read and hybrid *de novo* assemblies exemplary for the discovery of genomic variants and the prediction of genes.

The following background section provides basic information about the topic of *de novo* assembly in general and the Illumina and ONT sequencing technology. Thereafter, a top-level overview of the used materials and methods is given. After presenting and discussing the therewith obtained results, a conclusion regarding a best practice for state of the art *de novo* assembly is drawn. Supplementary material and a detailed description of the methods usage are accessible via the projects GitHub repositroy at https://github.com/Integrative-Transcriptomics/TechnicalReport-AssemblyBestPractice.

## Background

The following section summarizes relevant aspects of *de novo* assembly related to the addressed questions. A detailed theoretical and methodological definition of the individual aspects is not provided, but interested readers are referred to corresponding sources.

### Challenges of *de novo* assembly

The problem of *de novo* assembly is not solely set up by the computational hurdle of deriving a continuous genomic region in its correct order from shorter fragments, but also heavily influenced by a variety of problems that arise due to the underlying sequencing procedure. Herein three of these problems, namely (1) possible errors in the read sequences, (2) uneven or no coverage across the genome and (3) the problem of multiple consecutive copies of the same sequence within a genome, referred to as repeats, will be considered in more detail [2].

Given a DNA sequence as a string of the symbols A, C, G and T, representing the canonical DNA bases, sequencing errors can be classified as substitutions, deletions and insertions, visualized in figure 1. Substitutions are also referred to as mismatches, while positions at which the base symbols are equal are referred to as matches. Insertions and deletions are often collectively referred to as indels. To estimate sequencing errors, a to-date standard for all sequencing technologies is the Phred quality score as given in definition 1 [2].

**Figure 1:**
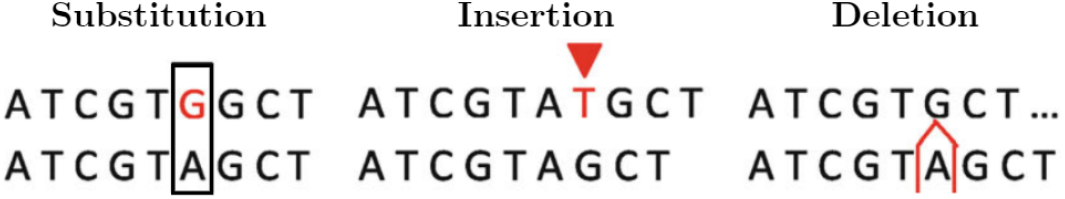
Visualization of sequencing error types. The error position in the upper sequence relative to the lower sequence is highlighted in red. The image is adopted from Liao *et al*., figure 4 [8].

#### Definition 1

(**Phred quality score**, adopted from Myers [2]). *The Phred quality score of a read position is given as*

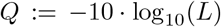

*where L is the likelihood that the base assigned to the position is incorrect*.

These quality scores are computed for each read position with machine learning techniques on features of the respective sequencing device [2].

A more generalized method to evaluate the performance of sequencing devices is the average read accuracy: Thereby the number of matches relative to a reference read is divided by the sum of match-, mismatch-, deletion- and insertion-positions for a given read [5]. However, this method may be influenced by the respective algorithm used for read alignment [5].

Sequence alignment in principle refers to the problem of inferring a similarity score of two or more sequences by matching as many positions with the same symbol as possible whereby gap symbols may be introduced to account for insertions and deletions [10].

This problem is not to be confused with sequence mapping, being of independent interest, with which it is tried to find the position of a string (e.g. a read) in a set of larger strings (i.e. one or multiple chromosomes) [10]. A comprehensive overview of both, sequence alignment and mapping, is given by Altschul and Pop and will not be covered in this report [10].

The second problem mentioned, irregularities of read coverage, can be considered in two variants: The number of times a specific position in the genome was sequenced redundantly, which is referred to as the *depth of coverage* (DOC).

The percentage of positions of a genome that have been sequenced at least *x* times, where *x* is a reasonable large enough natural number, referred to as the *breadth of coverage* (BOC) [11].

For *de novo* assembly the BOC should be that high that each genomic position was at least seen once, while a high DOC may be exploited to eradicate sequencing errors [11].

Lastly, the problem of reconstructing repeat regions is closely related to read length limitations: In order to reconstruct a repeat region, one read spanning the full genomic section in which the repeats occur has to be obtained [7].

### DNA sequencing technologies

An overview of the basic sequencing procedure of Illumina and ONT devices is given in figure 2.

**Figure 2:**
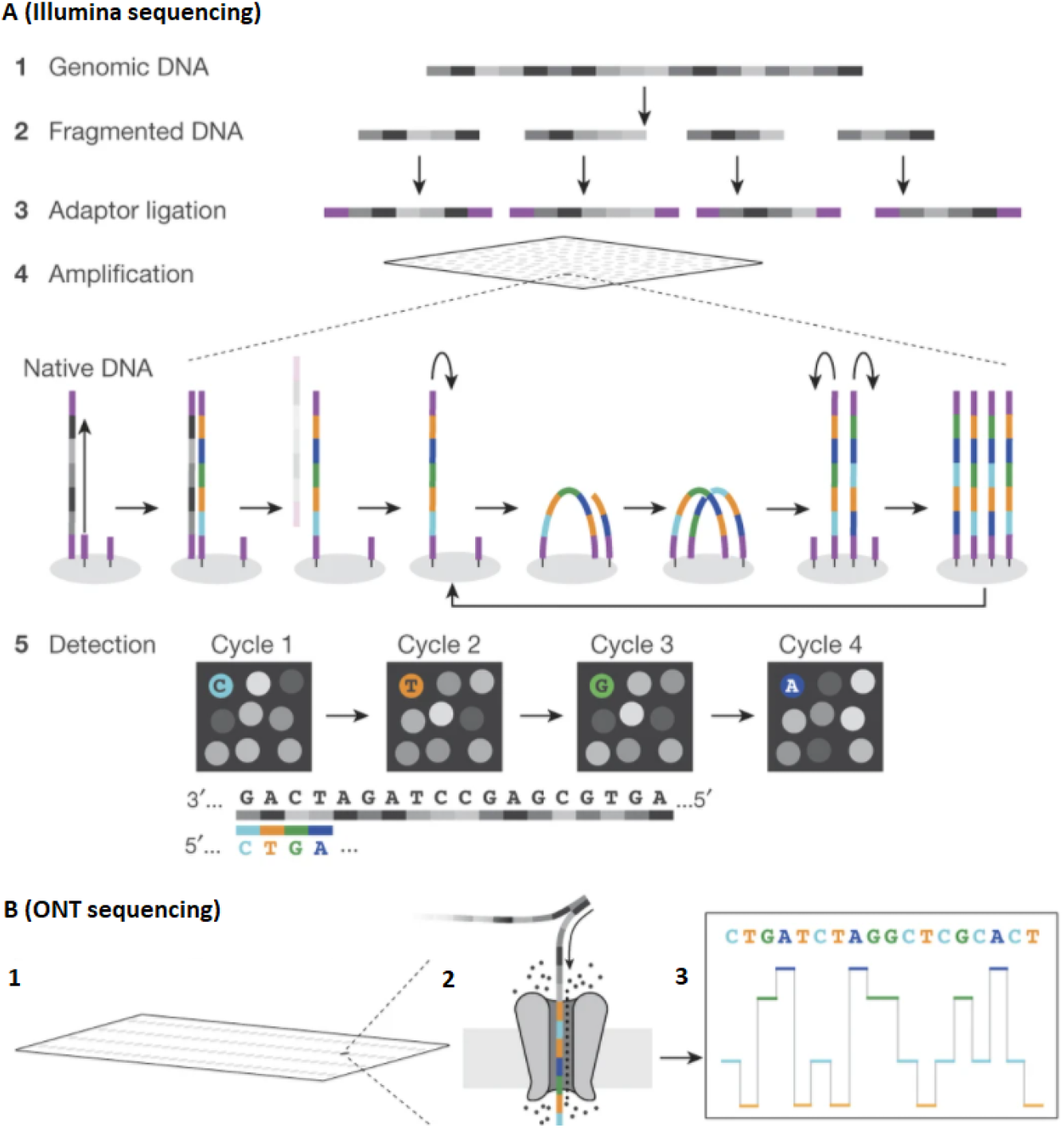
Schematics of the Illumina and ONT sequencing procedure. **(A, Illumina sequencing) 1-2** Sampled DNA is fragmented by enzymatic digestion or sonication. **3** The fragment ends are ligated with adapters. **4** The adapters hybridize with short nucleotide sequences mounted on a flow cell. Once hybridized, a bridge amplification process is repeatedly performed to create a cluster of copies of the same fragment. By removing DNA strands with a specific orientation, only strands with the same orientation remain in each cluster. **5** Recurringly, fluorescent labeled nucleotides are passed to the flow-cell and a device measures which nucleotide type was incorporate at each cluster site. The sequence of these light signals is used to infer read sequences [9]. **(B, ONT sequencing) 1** Devices comprise flow-cells spanned by a membrane in which protein nanopores are embedded and on which an electric potential is applied. **2** This potential causes ions and sample DNA to flow through the nanopores. Motor proteins unzip double stranded DNA and guide one strand through the pore. Thereby, the nucleotides inside the nanopore affect the ion flow in a distinct way and cause changes of the electric current. **3** This sequence of changes in the electric current is used to infer read sequences [5]. The image is adapted from Shendure *et al*., figure 1 [1].

Illumina sequencing devices (i.e. the Illumina MiSeq device is considered herein for a comparison) achieve a read accuracy of above 99 % at a per Gigabase cost of about 25 $ for a 55 h run and a read length of 300 base pairs [5].

The bridge amplification step is not only prone to substitution errors that will be contained and amplified in each cluster, but also to a base composition bias favoring regions with balanced GC base content [8]. The latter may results in an uneven depth of coverage across the genome [8]. However, the main error source is the asynchronous incorporation of nucleotides – also referred to as de-phasing – into the population of one cluster, leading to an increased noise of the fluorescence signals and thus an increased error rate as the length of the molecules increases [3]. This effectively limits the read length [3], what can be considered as the major drawback of this technology as they are lacking potential in resolving genomic repeat regions [9].

To overcome this problem protocols for paired-end sequencing can be used [9]. Thereby, both ends of the original DNA fragments are sequenced, yielding not only a higher data throughput but information about the distance between the two reads on the original fragment [9]. While this information can be used to reconstruct repeat regions, it does not suffice to resolve all of such [9].

In contrast, ONT sequencing devices (i.e. the ONT MinION device is considered herein for a comparison) achieve a comparatively lower read accuracy of about 85 % at a per gigabase cost of 35 $ for a 48 h run and mean read lengths of 50 · 10^3^ base pairs [5]. The read length is thereby dependent on the fragment size of the sampled DNA and in practice reads spanning a million base pairs were observed [5]. ONT reads are thus well suited for reconstructing repetitive sequences, as they are likely to span over the whole region in which the repeats occur [7].

High error rates are the biggest drawback of ONT technology: As a DNA strand passes through the nanopore, the structural similarity of the nucleotides and the fact that multiple nucleotides influence the current changes at once make it difficult to assign one distinct current value to each nucleotide type [5]. Furthermore, a non-uniform translocation speed of the DNA strand is hampering the identification of especially homo-polymers as for such no changes in the current are observed [5]. The latter gives rise to insertion and deletion errors which do not occur for second generation technologies [5].

Another possible error source is the process of translating the measured electric current changes into reads – referred to as basecalling [5]. The most recent basecallers, first published in 2017, are Albacore and Guppy – a GPU accelerated version of Albacore – which both rely on the same neural network architecture and achieve read accuracies of 87 to 88 % [12]. The process of basecalling will not be covered in this report but interested readers are referred to a review by Rang *et al*. [5] and a recent benchmarking of basecallers conducted by Wick *et al*. [12].

As these machine learning base callers need to be trained, but DNA samples show different characteristics regarding, for example, GC content, DNA modifications and codon usage, the nature of this training data sets may affect the basecalling performance [5].

### Algorithmic methods for *de novo* assembly

To date two major algorithmic frameworks are used to derive a contiguous genome from a set of reads. The fist framework is the *overlap layout consensus* (OLC) paradigm, which will be recapped based on the description of Myers [2]. Thereby, the assembly procedure is conducted in three steps. First, all suffix-prefix overlaps of any two reads of the given set of reads are computed. In order to obtain not too few but significant overlaps, that is to say to minimize the number of overlaps occurring by chance, the length of the overlap and the number of allowed errors in the string matching have to be adjusted. In a second step, an overlap graph is constructed by introducing one vertex for each read and by drawing an edge between any two vertices if there exists a presumably non-random overlap between the respective reads. Next, Hamiltonian paths are searched in this graph, mostly via greedy heuristics, each giving rise to a contiguous genomic sequence by concatenating the strings represented by each vertex along that path. Finally, a consensus sequence is computed from the sequences derived from these paths.

The second framework is the utilization of Eulerian *de Bruijn* graphs recapped herein based on the description of Liao *et al*. [8]. For such, a vertex is introduced for each distinct *k* − 1-mer (i.e. for each distinct sub string of length *k* − 1 of any read) and an edge is drawn between two vertices if the *k* − 1-mer represented by the source and sink vertex is a prefix and suffix, respectively, in any read. A contig can then be sought of this structure by walking along an Eulerian path.

Both of the described frameworks differ in their applicability for either ONT long reads or Illumina short reads [2]. The most striking issue regarding the OLC approach is the fact that short overlaps, as they may occur for short read lengths, will lead to a collapse of most of the repeat regions of the true genome [2].

In contrast, for the *de Bruijn* approach reads have to be highly accurate as contigs are formed only from adjacent *k* − 1-mers in reads an high error rates could result in many disconnected graph components and dead ends [8].

A more detailed description, including the computational complexity of the two frameworks, can be obtained from the respective reviews of Myers [2] and Li *et al*. [8].

However, none of the above mentioned frameworks were especially tailored for long, rather erroneous, reads [7]. To overcome this problem hybrid approaches combine short and long reads either directly for assembly or use accurate short read or high coverage long read information to correct sequencing errors before or after assembly – a process commonly referred to as polishing [7].

### Evaluating *de novo* assemblies

One commonly used approach to assess contig quality is to consider the contiguity, completeness, and correctness of the derived contigs relative to a reference genome [13]. A perfect genome assembly would be contiguous, complete and correct [13].

Contiguity basically reflects the fragmentation of the genome as the size and number of contigs and how these can be aligned to a reference genome without larger indels [13].

Completeness can be considered as the gene content of the contigs compared to the reference as well as the overall percentage of the reference genome that was reconstructed. Finally, correctness mainly concerns the ordering and location of contigs or sub-segments of such relative to the reference sequence as well as errors in the sequence [13]. Incorrect rearrangements may be classified as inversions, relocations and translocations, with respect to a true reference genome [13]. These are graphically represented in figure 3.

**Figure 3:**
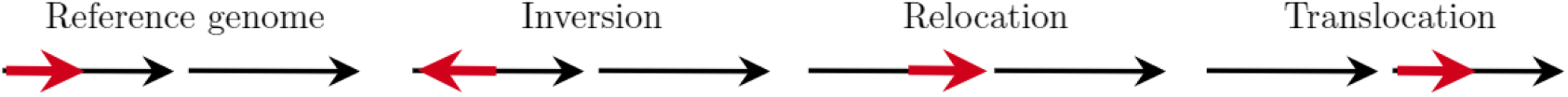
Graphical representation of genomic rearrangements. Each black arrow represents one genomic segment, for example a chromosome, and the arrowhead indicates its orientation. The red arrow represents the segment of interest. Its location and orientation on the right most sub-figure relative to the three remaining sub-figures describe the respective rearrangement. The three error types may also occur in combinations and switch strands. The image is based on descriptions from Salzberg *et al*. [14].

Obliviously, the three characteristics stated above can not be viewed as separated from each other, as for example genomic rearrangements and errors could lead to genes not being detected or prevent contigs from being joined together during the assembly process.

Furthermore, it has to be thought of the fact that such errors may not only arise due to the assembly software but may also originate from errors of the sequencing process or may even be true changes in the genome of the sampled organism relative to the reference sequence.

Once an assembly of sufficient quality is completed it may be used in a variety of downstream applications, for example the prediction of genes or discovery of nucleotide variations.

### Genetic variant discovery

The discovery of genetic variants aims at finding *single nucleotide variants* (SNVs), i.e. substitutions, insertions and deletions of single nucleotides relative to a reference genome, large genomic rearrangements but also gene or plasmid copy number variations [7].

In this report it is focused on the discovery of SNVs in form of single base substitution. For this task one can either use the sequence derived from a trustworthy *de novo* assembly and align it against a reference genome to detect positions with differing bases or directly map and align the input reads to a reference genome [7].

The latter approach can yield additional confidence information, as thereby the DOC and read Phred quality scores can be taken into account to compute likelihoods for the reads supporting an alternative or the reference allele [7]. This information is commonly stored in so called vcf files which contain not only information about the position, reference and alternative base but also the mentioned likelihoods and statistics regarding how many reads cover the respective position as well as the reads quality [15]. A comprehensive overview about computing the mentioned likelihoods is given by Li [16].

## Material and Methods

### Assemblers and polishing tools

For selecting promising assemblers, recently published reviews were examined. Based on a benchmark of long read assemblers on prokaryote data sets from Wick and Holt [6] the following long read assemblers were selected:

- **Canu** exploits the OLC paradigm as assembly strategy and additionally conducts a prior long read correction and trimming [17].
- **Flye** uses a structure similar to *de Bruijn* graphs, so called repeat graphs, which allows approximate sequence matches during construction as well as an integrated polishing step [18].
- **Raven** uses the OLC paradigm for assembly and polishes the derived contigs with Racon [19].

Based on a benchmarking of hybrid assemblers from Chen *et al*. [20] and a study from Sydenham *et al*. in which different assemblers were compared in their ability to compute *Bacteroides fragilis de novo* assemblies [21], the tool Unicycler was included as hybrid and long read assembler. Furthermore, the recently published hybrid assembler HASLR [**?**] was selected.

- **Unicycler** uses the OLC paradigm for long read input. The derived contigs are polished with Racon in advance. For hybrid input, a *de Bruijn* graph is built from short reads. Next, long reads are used to derive unambiguous paths in this graph and the resulting contigs are polished with Pilon [22].
- **HASLR** first constructs short read contigs which are then mapped to long reads. From this information a backbone graph is built of which the final assembly is derived [23].

Furthermore, the assembly consensus tool **Trycycler** was selected [24]. It receives a set of contigs from multiple assemblies as input and proceeds in six steps, mainly comprising a similarity based clustering of the input contigs, an orientation correction and circularization as well as a final consensus computation via multiple sequence alignment [24].

As for polishing tools the following three were relevant for this project as they were either used as external tool or internally by one of the selected assemblers:

- **Medaka** is an assembly polishing tool developed by ONT which performs polishing with long reads using neural networks [25].
- **Pilon** is used to identify errors in draft assemblies from a mapping of short reads to a draft assembly [26].
- **Racon** maps and aligns reads (long or short reads) to a draft assembly and conducts polishing by computing a multiple sequence alignment via so called *partial order alignment* graphs [27].

### Acquisition and quality control of read data

For this project it was decided to collect real prokaryotic whole genome sequencing data. These reference data sets comprise two publicly available read sets from a study of De Maio *et al*. [28] and one non-public read set from the collaborative research centers Transregio 261 of the Deutsche Forschungsgemeinschaft.

The publicly available sets comprise whole genome sequencing data of *Escherichia coli* CFT073 (BioSample accession SAMN10819847) and *Klebsiella pneumoniae* MGH78578 (BioSample accession SAMN10819805) isolates [28]. Both comprise paired-end read sets with a maximal read length of 150 base pairs generated with the Illumina HiSeq 4000 platform and long read sets generated with the ONT MinION device and basecalled with Albacore version 2.0.2 [28]. In the following the sets will be addressed with their respective strain identifiers CFT073 and MGH78578.

The read set from the Transregio 261 comprises whole genome sequencing data of a *Staphylococcus aureus* RN4220 strain isolate. It comprises one Illumina paired-end short read set, generated with the Illumina MiSeq device and a maximal read length of 150 base pairs, and two ONT long read sets, sequenced with an MinION device and basecalled twice with Guppy versions 3.2.1.0 and 4.0.1.1. In the following this set will be addressed with its strain identifier RN4220.

The following reference genomes and associated gene annotations, which were specified by De Maio *et al*. for the public data sets [28], were collected for evaluating the computed assemblies:

- For **CFT073** a complete chromosome sequence spanning 5, 231, 428 base pairs provided by Welch *et al*. with 5, 288 annotated features (RefSeq accession NC 004431.1) [29].
- For **MGH78578** a complete chromosome sequence spanning 5, 315, 120 base pairs and five complete plasmid sequences spanning 175, 879, 107, 576, 88, 582, 4, 259 and 3, 478 base pairs, respectively, provided by McClelland *et al*. and with 5, 689 annotated features (RefSeq accession NC 009648.1 to NC 009653.1) [30].
- For **RN4220** a genome fragmented into 118 contigs with a total length of 2, 663, 395 base pairs provided by Nair *et al*. and annotated with 2, 913 features (RefSeq accession NZ AF-GU00000000.1) [31].

The acquired long reads were further processed with Porechop to trim remaining adapters from the ends or the middle of the reads [32].

After this step all read sets were subject to a quality control step with FastQC [33].

Additionally, the DOC and BOC of the read sets with respect to the selected reference genomes was assessed by first mapping the reads to the reference genomes with minimap2. minimap2 follows a seed-chain approach for mapping and was especially designed for both, accurate short reads and more erroneous long reads [34]. Thereby, parameters were set to account for the read type and in order to output files in SAM format. The resulting SAM files were subject to sorting and conversion to BAM files using samtools [35]. Finally, these BAM files were processed with the samtools depth command in order to count the number of aligned reads per reference position.

From the therewith obtained counts the mean DOC and BOC (considering positions with at least one aligned read) was computed. The BOC was compared with the formula given in definition 2 to validate the underlying sequencing procedures of the read sets.

#### Definition 2

(**Expected BOC**, based on Lander and Waterman [36]). *By considering the number of times a position being sequenced as a random variable following a Poisson distribution*

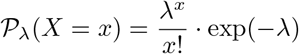

*where the parameter λ is the expected DOC, the expected BOC, with respect to each position being sequenced at least once and given a mean DOC of c, is given as*

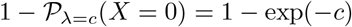

In order to assess the DOC not only as the mean value of the full reference set, but in a higher resolution along the reference genome, the per position DOC was visually explored.

### Read data sub sampling

To assess the influence of low long read coverages to the assembly quality, sub samples of the long read sets had to be genereated. For this it was decided to generate sub samples of 200, 150, 100, 80, 60, 40, 20, 15, 10, 8, 6, 4, 2 and 1 mean DOC of the CFT073 and RN4220 long reads.

This was done with Rasusa [37], a tool which computes the number of base pairs needed to achieve an expected depth of coverage relative to a reference genome length and randomly samples reads until this number is reached [37].

For the approximate genome lengths the length of the CFT073 reference genome was rounded off to 5.2 *·* 10^6^ base pairs.

For the RN4220 reference genome it was decided to use 2.8 *·* 10^6^ base pairs, the median length of 12, 401 genome assemblies of *S. aureus*, due to the high fragmentation of the reference [38].

To check if the expected coverage is met, the before described procedure of DOC and BOC quality control was applied to each read sub sample. With the therewith obtained results the sub sampling was validated by computing the (non absolute) difference in means for the expected DOC, i.e. the one specified for Rasusa, and the true DOC as well as the expected BOC, based on definition 2 by using the true mean DOC as parameter, and the true BOC.

### Realisation of assemblies

The selected assemblers were run, if possible with default parameters, on all reference sets as well as on the CFT073 sub samples, i.e. for each assembler four assemblies with the full data sets and 14 assemblies with sub sampled data were computed. Regarding the RN4220 sub samples it was decided to only use the hybrid assembler Unicycler to assess the effect of an improved basecaller version. As Canu does not include an internal polishing step, what is the case for the other assemblers, the Canu assemblies were additionally polished using Medaka.

Furthermore, Trycycler was run on all four long read assemblies, whereby the polished Canu assembly was chosen if possible, of the three reference sets and the sub samples of the CFT073 read set. Each computed Trycycler assembly was additionally subject to polishing with Medaka.

To assess the quality difference of approaches incorporating short reads directly into the assembly process and the pipeline of using short reads only for polishing subsequently of the actual assembly process, the Trycycler consensus assemblies were additionally polished using Pilon.

### Assessment of assembly quality and genetic feature prediction

All of the created assemblies were evaluated using Quast [39]. Quast receives an assembly and one or multiple of the reads used to compute the assembly, a reference genome and gene annotations of the reference genome [40]. Quast can optionally be used to run GeneMark (i.e. GeneMarkS), a tool for the prediction of genetic features based on Hidden Markov Models [41]. In what follows a brief description of the herein considered metrics as they are defined in the Quast manual and their relation to the aspects completeness, contiguity and correctness as described in the background section is given [40]:

The assembly completeness was assessed based on the genome fraction, that is the number of positions that were aligned from the assembly to the reference genome relative to the total number of reference positions, and the number of completely recovered genomic features divided by the number of annotated reference features.

The contiguity of the assemblies was assessed based on the number of contigs and the LA75 value, which is defined as the number of misassembly free continuously alignable contig sub segments of the assemblies to the reference genome that constitute for 75 % of the number of reference genome positions.

Finally, the correctness of the assemblies was assessed by the number of mismatches and indels per 100, 000 base pairs (100 kbp) of the reference genome and the number of misassemblies.

The latter is separated into four classes: Relocations occur if a contiguous part of a contig has to be split into two sub contigs for alignment to the reference genome and one of these sub contigs aligns at least 1, 000 base pairs away from the other one. Inversions and translocations occur in the same manner, but if one sub contig aligns on the reverse strand without positional shift or to another chromosome, respectively. In addition local misassemblies are defined as relocations but with a shift between 65 and 1, 000 base pairs.

We decided to run Quast with the reference genome and its annotation, run gene prediction and disable the discovery of nucleotide variations. Apart from this, default parameters were chosen, i.e. indels are reported up to a length of 65 base pairs and contigs are reported up to a length of 500 base pairs.

The gene prediction was run in order to assess qualitative differences regarding this of long read and hybrid assemblers.

### Discovery of single nucleotide variations

To investigate the potential of combining long and short reads for the discovery of SNVs the following procedures were applied:

Firstly, a mapping based approach was run with bcftools using either long reads, short reads or both read sets of the CFT073 reference, thus yielding three SNV sets. bcftools reports a quality score for each SNV as the Phred scaled likelihood of the assertion of an alternative base being wrong, i.e. high numbers indicate a high confidence, and a Phred scaled genotype likelihood rounded to the closest interger, i.e. high numbers indicate a unlikely genotype, for every genotype per sample [15]. For the mapping, the same procedure as for the coverage analysis was applied. The resulting files of the mapping were further processed using the bcftools mpielup and bcftools call commands to generate a vcf file.

Thereby, a ploidy of one (as the CFT073 reference without plasmids can be considered as a haploid organism) was specified, thus genotypes are considered as reference or alternative, and the detection of indels was disabled.

Secondly, an alignment based approach was run by using Mauve [42] and DNAdiff, a wrapper for MUMmer [43]. Both tools basically conduct a whole genome alignment and report differing bases as SNVs. The long read consensus assembly derived with Trycycler polished with Medaka and the hybrid assembly derived with Unicycler were subject to the two methods, as these were expected to be of the highest quality among their assembler classes, yielding a total of four SNV sets.

Prior to any downstream analysis of the detected variations it was necessary to specify a set of high-confidence reference SNVs. For this a consensus approach was chosen in which SNVs were included into the set of reference SNVs if any two of the following critera agreed on the position and alternative base of the SNV: (1) The SNV occurs in the SNV set derived with bcftools on the short reads, has a DOC of at least 40 and a Phred scaled quality score of at least 50, i.e. the likelihood that the assertion is wrong equals 10^−5^. (2) The SNV occurs in the SNV set derived with DNAdiff from the Unicycler hybrid assembly. (3) The SNV occurs in the SNV set derived with Mauve from the Unicycler hybrid assembly.

By including the reference SNV set, the pairwise intersection of all SNV sets was computed.

Finally, it was evaluated to which extend the SNVs detected with bcftools, using short reads, long reads or both sets, agreed upon the reference and non-reference SNVs by comparing the reported per sample genotype likelihoods and alternative base calls.

## Results

### Assessment of read quality

A summary of the results of the long read processing with Porechop and quality control step of both, short- and long reads, with FastQC is given in the following. Regarding Porechop, start adapters were removed from 95.9 %, 94.6 %, 98.8 % and 98.8 % of the reads and end adapters were removed from 60.2 %, 64.9 %, 59.1 % and 64.7 % of the reads while 37, 19, 224 and 257 reads were split due to middle adapters, for the CFT073, MGH78578, RN4220 (Guppy 3.2.1.0) and RN4220 (Guppy 4.0.1.1) long read sets, respectively. All following results were obtained with the trimmed long reads.

Concerning the FastQC reports, a warning or failure was given regarding the reads quality scores and length distribution for all long read sets. A length distribution warning was given for all short reads as well, revealing that a few hundreds to thousands of reads had a length shorter than 150 base pairs.

Furthermore, failures or warnings were given for the GC content, deviations from the expected per position base distribution and *k*-mer content for all read sets. Thereby, the latter two concerned the beginning of the reads exclusively, that is to say up to the tenth read position, and the observed GC content distributions correspond to the expected distributions, but with a lower variance leading to a more narrow peak around the references GC content. None of the short reads showed remaining adapters, however the RN4220 short reads showed a ten- to fifty-fold duplication level for about 25 % of the read sequences.

An overview of the reads quality, mean DOC and BOC is given in table 1.

**Table 1:**
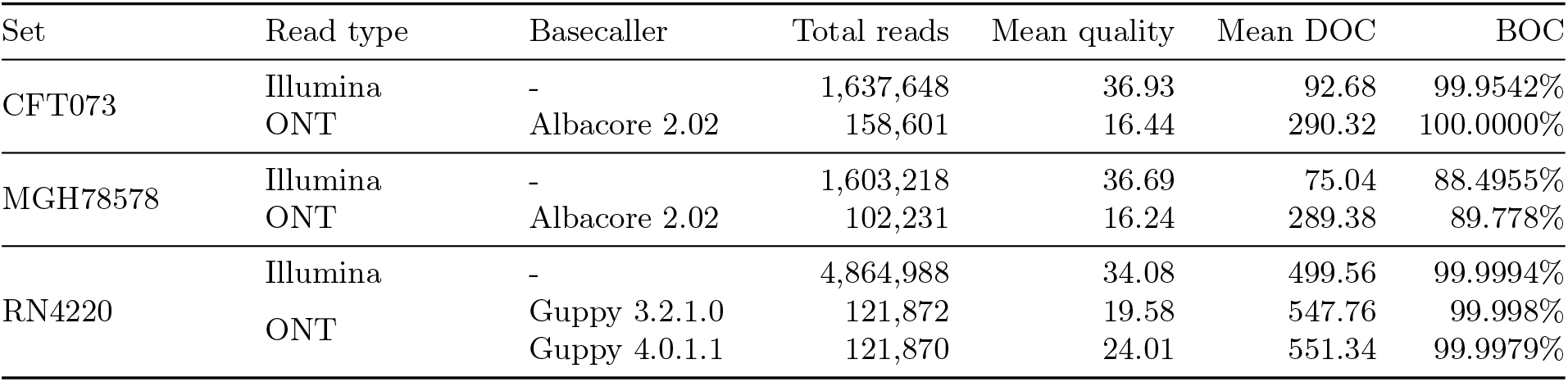
Overview of read quality and coverage statistics. ONT rows concern the trimmed long reads. The total number of reads and mean quality, i.e. the mean Phred quality score of the read set, were reported with FastQC. The mean DOC and BOC were computed based on the results of mapping with minimap2 and subsequent analysis with samtools depth.

While the mean read quality of long reads appeared to be higher for more recent basecaller versions, they still did not reach the ones of the Illumina short reads. Furthermore, all sets showed high DOC mean values, each giving rise to an expected BOC of 100 % according to definition 2. By considering this it was striking that the BOC of the CFT073 short reads and all three read sets of RN4220 appeared to be slightly lower than 99.9 % whilst for the MGH78578 reads more than 10 % of the reference genome was uncovered.

The investigation of the per position DOC revealed that the MGH78578 chromosome contained several areas of zero coverage and the five plasmids feature only few regions with non-zero coverage. The CFT073 and RN4220 sets showed a nearly uniform mean DOC.

Regarding the validation of read sub sampling, the difference in means was computed to be 0.174 for the DOC (wrt. the expected DOC defined for Rasusa) and −0.011 for the BOC (wrt. to the expected BOC computed with definition 2 given the true mean DOC) using all 42 read sub samples. Thus, on average the sub samples showed a slightly lower DOC than defined but a higher BOC than expected. For all following analysis the true mean DOC of the sub samples was used.

### Assessment of assembly quality

Hereafter, the completion of the assemblies will be described followed by presentation of the assembly quality characteristics computed with Quast.

All assemblers except HASLR were able to complete the assemblies on all four data sets. HASLR returned empty fasta files for all CFT073 and both RN4220 sets. For the MGH78578 set, HASLR was able to compute an assembly of 38, 398 base pairs length, but with a mismatch rate of 1, 890, 02 per 100 kbp. This assembly was removed for all subsequent analysis.

The Trycycler assemblies of the CFT073, MGH-78578 and RN4220 (Guppy 4.0.1.1) sets showed circularization issues of the Canu assemblies which were polished with Medaka and therefore the unpolished Canu assemblies were used. Trycycler was unable to circularize both Canu assemblies for the RN4220 (Guppy 3.2.1.0) set for which reason it was excluded. Furthermore, two contigs of the Unicycler and Canu assembly had to be removed for the MGH78578 set.

Regarding the completion of the CFT073 sub sample assemblies, Trycycler was unable to conduct all steps for mean DOCs below 37.58. Raven was unable to complete assemblies for sub samples with a mean DOC of 1.86 or lower and for the sub sample with a mean DOC of 1.86 Unicycler was unable to produce an assembly based on long reads only.

The reported metrics to assess the assemblies completeness, contiguity and correctness are visualized in figure 4.

**Figure 4:**
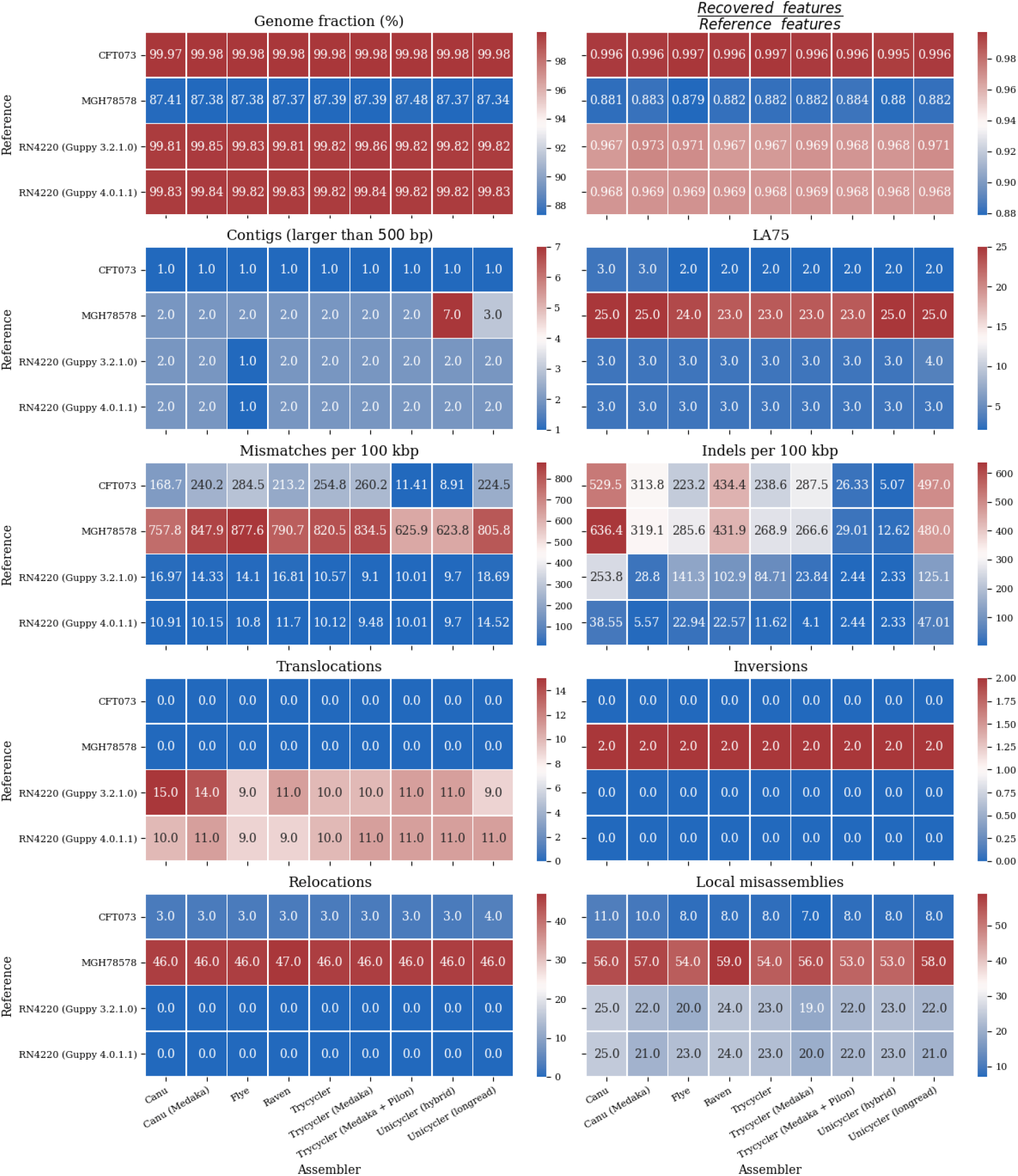
Assembly quality metrics reported with Quast. Each of the sub plots depicts one quality metric. The respective assembler and reference set of the assembly is denoted on the x- and y-axis, respectively. The axis tick labels are shared across sub plots. The coloring indicates the relative quality of the individual values: For the genome fraction (top, left) a high value is favourable while for the relative recovered genomic features (top, right) a value of 1 is optimal. The number of recovered features is reported wrt. completely recovered features. For all lower sub plots a low value is favourable. The number of mismatches and indels were computed per 100 kbp of the assembly being aligned to the reference genome.

It emerged that within each reference set the computed assemblies were equally complete. The same was observed for the contiguity, with the following exceptions: Unicycler was the only assembler that managed to reconstruct fragments of the MGH78578 plasmids yielding a total of 12 (of which 7 featured a length of 500 base pairs or more) and 3 contigs on hybrid and long read data, respectively. The alignment of these fragments to the reference sequence was assessed using Mauve. Furthermore, Flye was the only assembler able to assemble only one contig for both RN4220 read sets.

Concerning the correctness the Canu assemblies showed a tendency towards a higher number of local misassemblies compared to the other assemblers.

Besides that, the reported number of misassemblies within a set were also conserved. Striking differences were observed for the number of mismatches and indels.

Thereby, not only a separation into long read and hybrid assemblers became evident, whereas the hybrid assemblers scored better, but also a separation within the class of long read assemblers was observed: While the OLC based assemblers (Canu, Raven and Unicycler) were reported to have fewer mismatches as the *de Bruijn* based assemblers, the contrary was observed for the number of indels with an about two-fold increased number of indels of the OLC based assemblers.

In terms of correctness, the effect of polishing with Medaka and Pilon had opposing effects on the assembly correctness. While in the majority of the use cases the number of mismatches and indels was decreased, the contrary was observed too.

Except for the number of local misassemblies, the same metrics as used for the full read sets are visualized for the sub sample assemblies of the RN4220 reference in figure 5 and for the CFT073 reference in figure 6.

**Figure 5:**
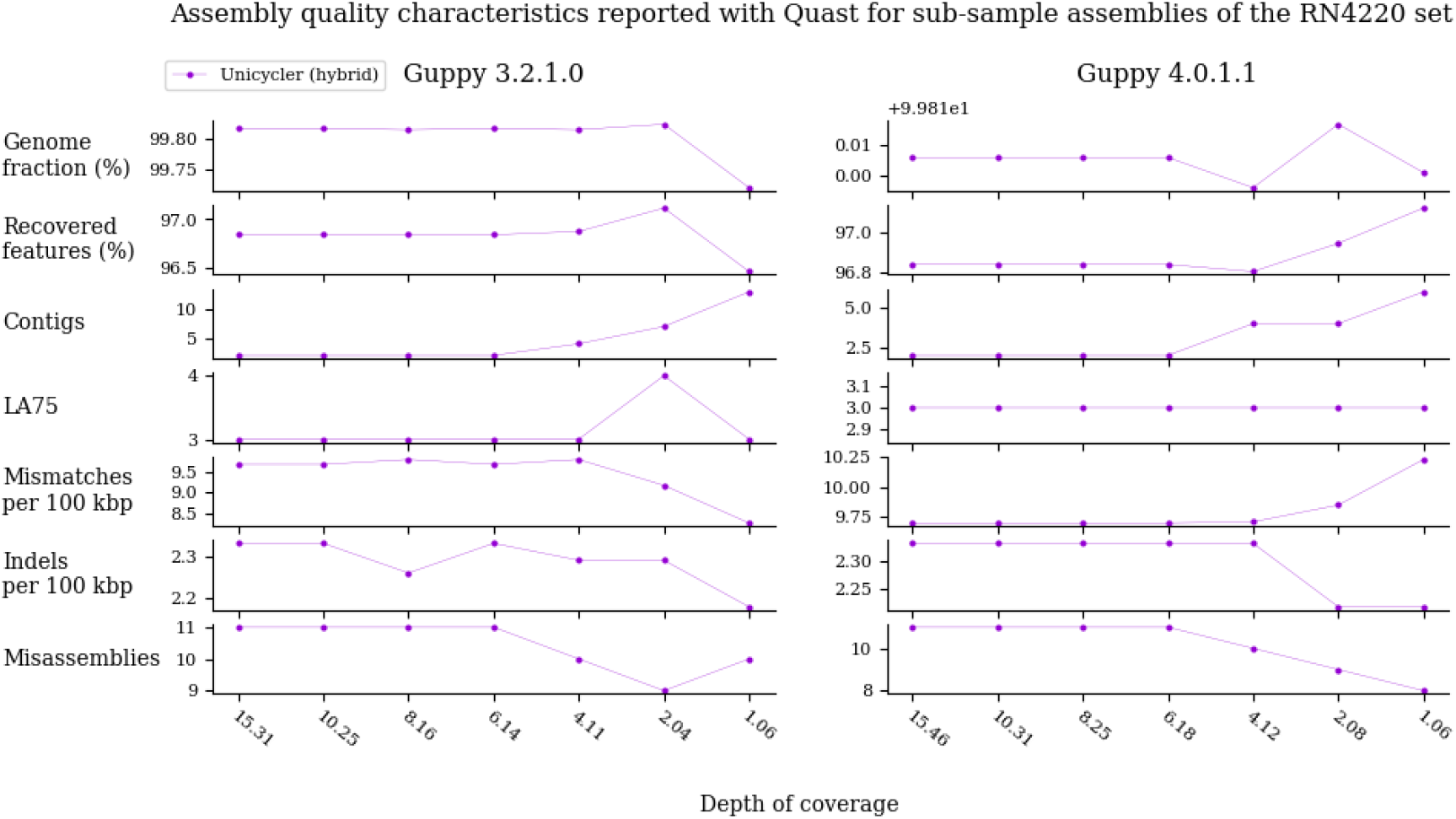
Assembly quality metrics reported with Quast regarding sub sample assemblies of the RN4220 set. Each row specifies one metric. The depths of coverage indicated on the x axis are the true mean values computed during quality control and shared among each row. Each dot corresponds to the value of the given metric computed for the assembler as indicated by the coloring in the legend. The number of misassemblies is the sum of reported translocations, relocations and inversions.

**Figure 6:**
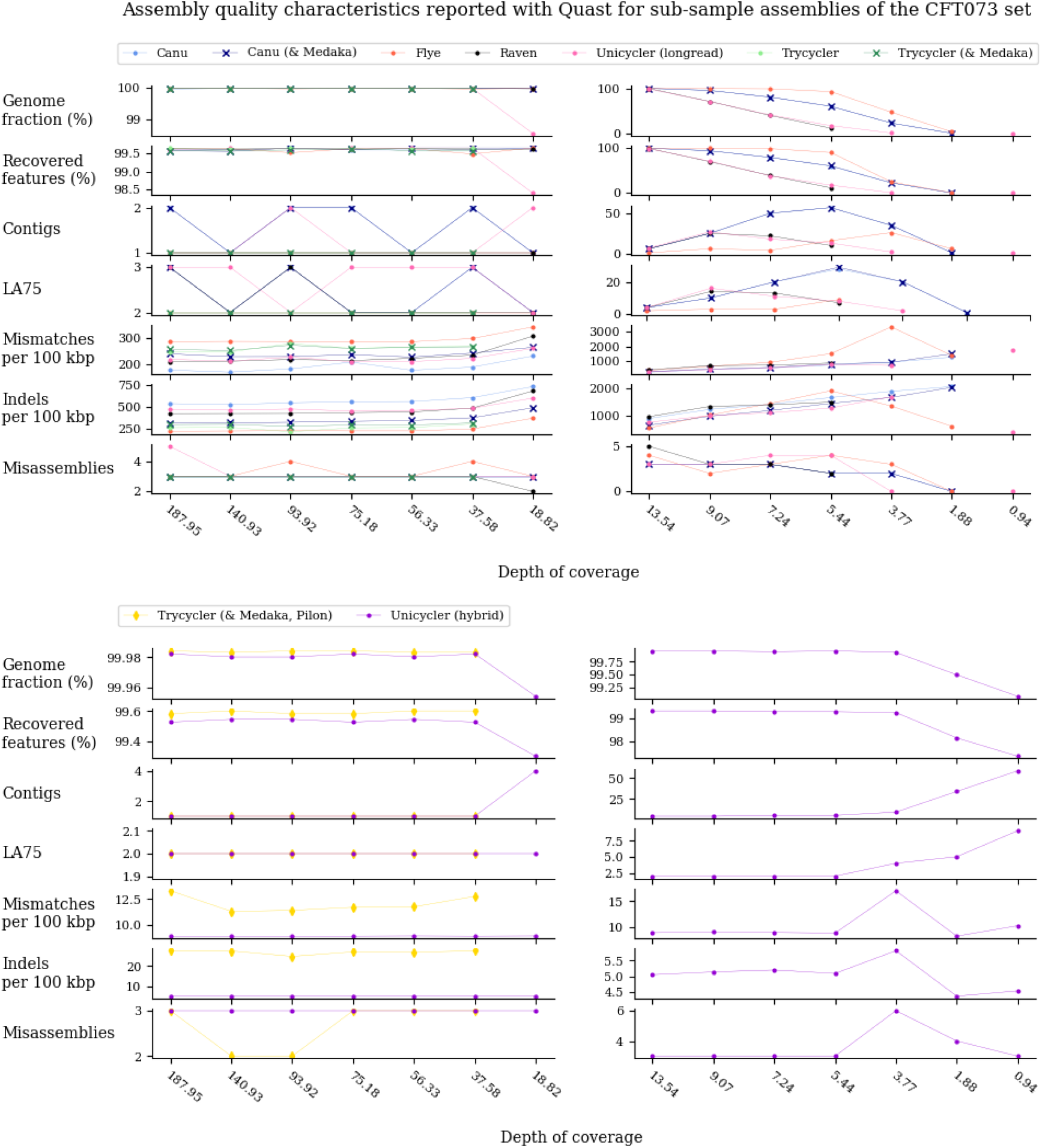
Selected quality characteristics reported with Quast of sub sample assemblies of the CFT073 set. The upper part of the plot depicts the values of the long read assemblers while the lower part depicts the values of the hybrid approaches. Each row specifies one characteristic. The depths of coverage indicated on the x axis are the true mean values computed during quality control and shared among each row. To achieve a higher resolution the plot separates assemblies of high coverage and such of low coverage sub samples into the left and right side, respectively. Each dot corresponds to the value of the given metric computed for the assembler as indicated by the coloring in the legend. The number of misassemblies is the sum of reported translocations, relocations and inversions.

For the hybrid Unicycler assemblies computed for the RN4220 sub samples no changes in the reported values were evident for mean DOCs above or equal to 8.16 for the Guppy 3.2.1.0 long reads and 6.18 for the Guppy 4.0.1.1 long reads, respectively. For lowering coverages the same pattern was observed as the number of contigs and accordingly the LA75 value increased, thereby this effect was pronounced stronger for the Guppy 3.2.1.0. reads as for the Guppy 4.0.1.1 reads. It was notable that the correctness regarding indels and mismatches of the assemblies increased for lower coverages compared to such sub sampled with a higher long read coverage.

Regarding the hybrid assemblies of sub samples of the CFT073 reference, the same pattern but with changes already occurring at a mean DOC of 37.58 and being more pronounced was observed.

For the long read assemblies of sub samples of the CFT073 set the reported values were observed to stay more or less stable for a mean DOC of above 56.33, while Canu and Unicycler yielded an oscilating behaviour for the number of contigs and the LA75 value.

For sub samples with a lower mean DOC all assemblers featured a worsening regarding completeness, correctness and contiguity. Regarding completeness, Flye was most resistant for lower mean DOCs followed by Canu, Unicycler and Raven.

The number of contigs and the LA75 values were observed to decrease again from a low enough mean DOC, being below 9.07 for Unicycler and Raven, below 5.44 for Canu and below 3.77 for Flye. These breakpoints appear to be related to low genome fraction of about 50 to 70 %.

Concerning correctness, the number of misassembly events gradually decreased while mismatches and indels increased with lower genome fractions for all assemblers. The majority of the assemblers stayed on their rank compared to the full data set, except for Flye being revealed to show comparatively high indel values.

### Assessment of predicted genetic features

The number of predicted genetic features reported with GeneMarkS are visualized, relative to the number of total annotated genetic features, in figure 7. It was striking that for the long read assemblies of the CFT073 and MGH78578 sets about one and a half as many genetic features were predicted as were annotated.

**Figure 7:**
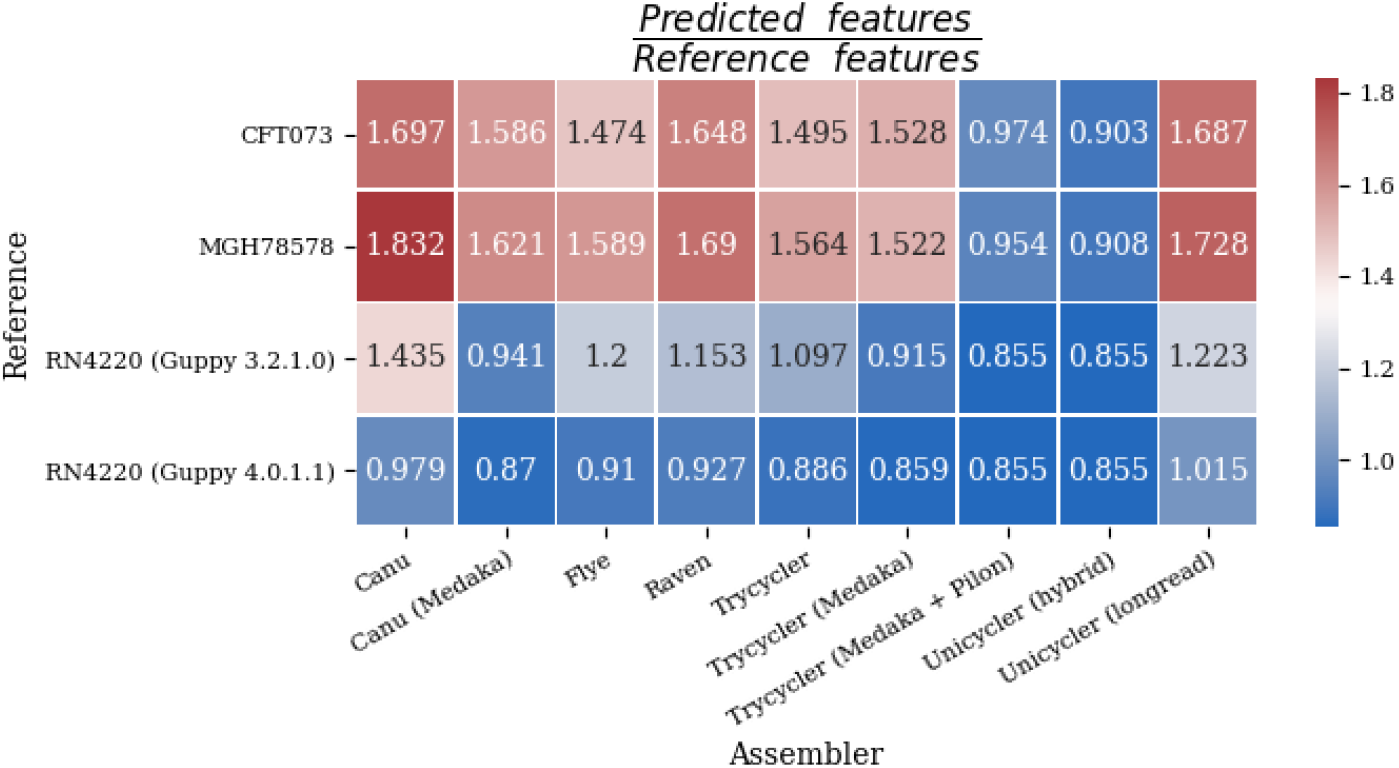
Overview of the number of predicted genetic features divided by the number of annotated genetic features for each set and assembler. The respective assembler and reference set of the assembly is denoted on the x- and y-axis, respectively. The coloring indicates the relative quality of the individual values: Here, a value of 1 is optimal. The number of predicted features is reported wrt. uniquely predicted features.

To assess this conspicuity the number of predicted and annotated features were compared for each assembly of the CFT073, MGH78578 and RN-4220 (Guppy 4.0.1.1) sets by grouping the annotated features by the number of predicted features that were contained to at least 60 % within the annotated feature and by comparing the length distribution of the annotated and predicted features. Therewith it was found out that the number of annotated reference features containing one or more predicted genes was higher for the long read assemblies of the CFT073 and MGH78578 set than for all hybrid assemblies and the long read assemblies of the RN4220 (Guppy 4.0.1.1) set. The distribution of feature lengths for the latter group matched the one of the annotated features, while this was not evident for the first mentioned assemblies. Furthermore, 200 to 600 less non overlapping features were counted for the hybrid assemblies of the CFT073 and MGH78578 set compared to the long read assemblies while this was vice versa for the investigated RN4220 assemblies.

### Assessment of discovered SNV sets

Finally, it was investigated to which extend long read and hybrid approaches yield different results for the discovery if single nucleotide variations in the form of single base substitutions.

The (in total 7) obtained sets, were analysed relative to each other by computing the size of the pairwise intersection of each of the sets. Each set was thereby considered as a set of tuples describing the position of the alternative base in the reference genome and the alternative base itself.

In addition a consensus set of the hybrid assembly alignment and short read mapping approaches was computed and used as reference SNVs. The therewith obtained counts are visualized in figure 8.

**Figure 8:**
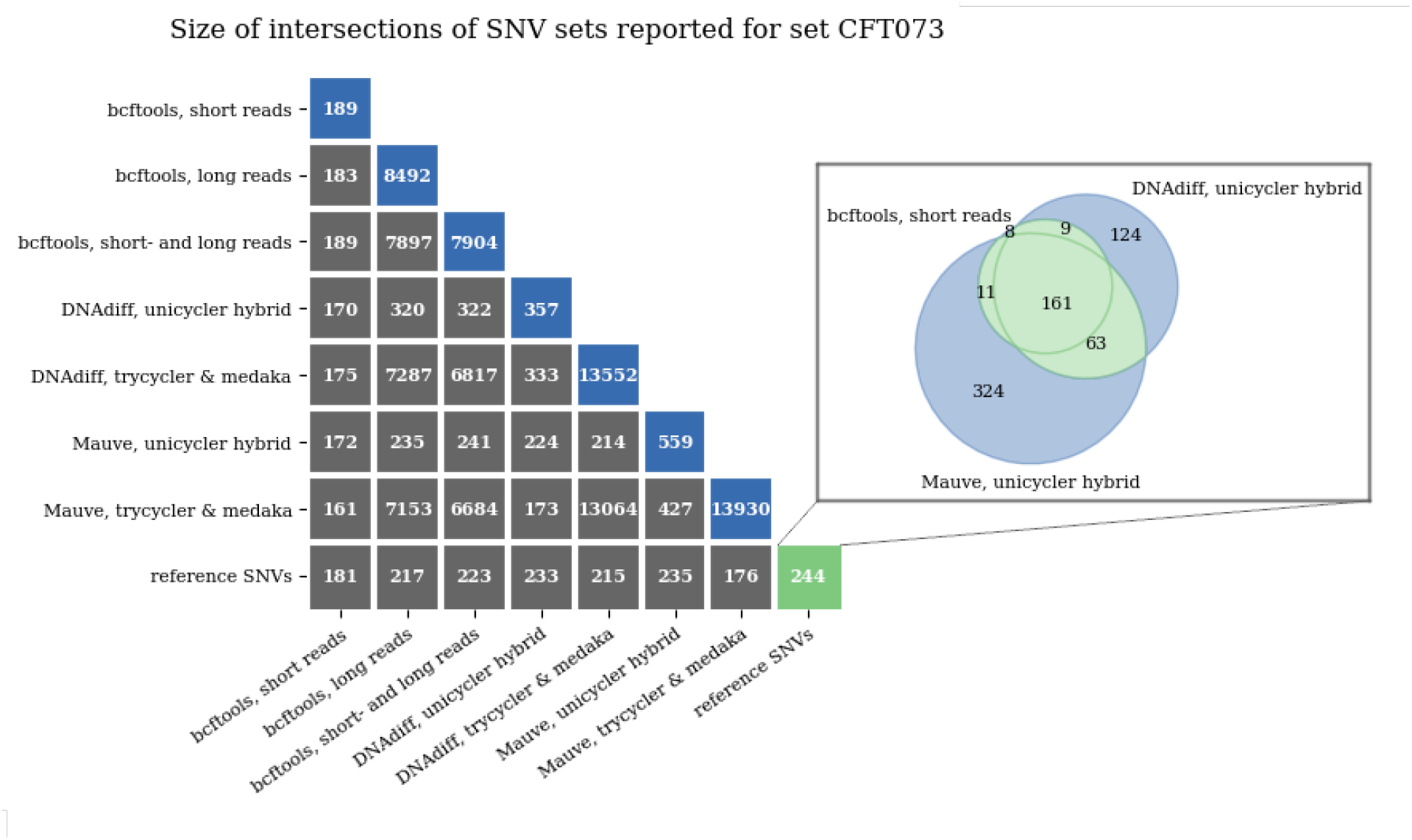
Pairwise intersection of SNV sets. The left side of the plot depicts the sizes of the pairwise intersections of all obtained SNV sets in gray and the size of the single SNV sets, i.e. the intersection of a set with itself, in blue. The last row includes a set of reference SNVs in green. The right side depicts how the reference set was computed.

It was ascertained that the approaches which used long read only assemblies or included long reads for the mapping based approach resulted in SNV sets that were many times larger compared to hybrid or short read only approaches. Furthermore, the size of the sets being based on assemblies are approximately the number of total mismatches relative to the reference genomes.

By considering the set of reference SNVs, containing a total of 244 positions, as positives in terms of a classification problem the long read approaches yield comparatively high numbers of false positives. This observation raised the question of if the long read approaches yield a high number of mistakenly asserted alternative base calls or if the usage of short read mapping and hybrid assemblies missed a high number of actually correct positions. To shed light on this question the with bcftools reported likelihoods for using short reads, long reads or both read sets, whereas for the third report these were reported with respect to both input sets, were analysed relative to the set of reference SNVs to assess the influence of the read types to each other. Regarding the set of reference SNVs the following six groups of genotype patterns were detected, whereas in no case a dissent about the alternative base was observed: 20 and 175 SNVs were called with no and with all bcftool runs, respectively.

23 SNVs were called by using long reads only and with respect to the long reads of the bcftools run with both read sets. 4 SNVs were called by using short reads only and with respect to the short reads of the bcftools run with both read sets.

20 SNVs were exclusively not called by using only short reads and 2 SNVs were exclusively not called by using only long reads.

Thus, for the third and fourth group the two read sets had contradicting likelihoods for an alternative base call, but one read types genotype likelihood out-weighted the other one. For the fifth and sixth group the genotype likelihood of one read set influenced the genotype call of the other set when using both sets – this pattern was observed if the confidence with respect to one read type was too low to yield an alternative base call.

For those SNVs that were called with bcftools using both read sets but not included in the set of reference SNVs the genotype likelihoods reported with respect to the short reads and long reads were compared by grouping the number of SNVs based on the difference of genotype likelihoods of the reference and alternative base call.

The results are visualized in figure 9 and indicate, that nearly all of the respective SNVs had a contrary genotype likelihood regarding the short and long reads. Thereby the short reads called the reference genotype with high confidence while the alternative genotype was called by the long reads.

**Figure 9:**
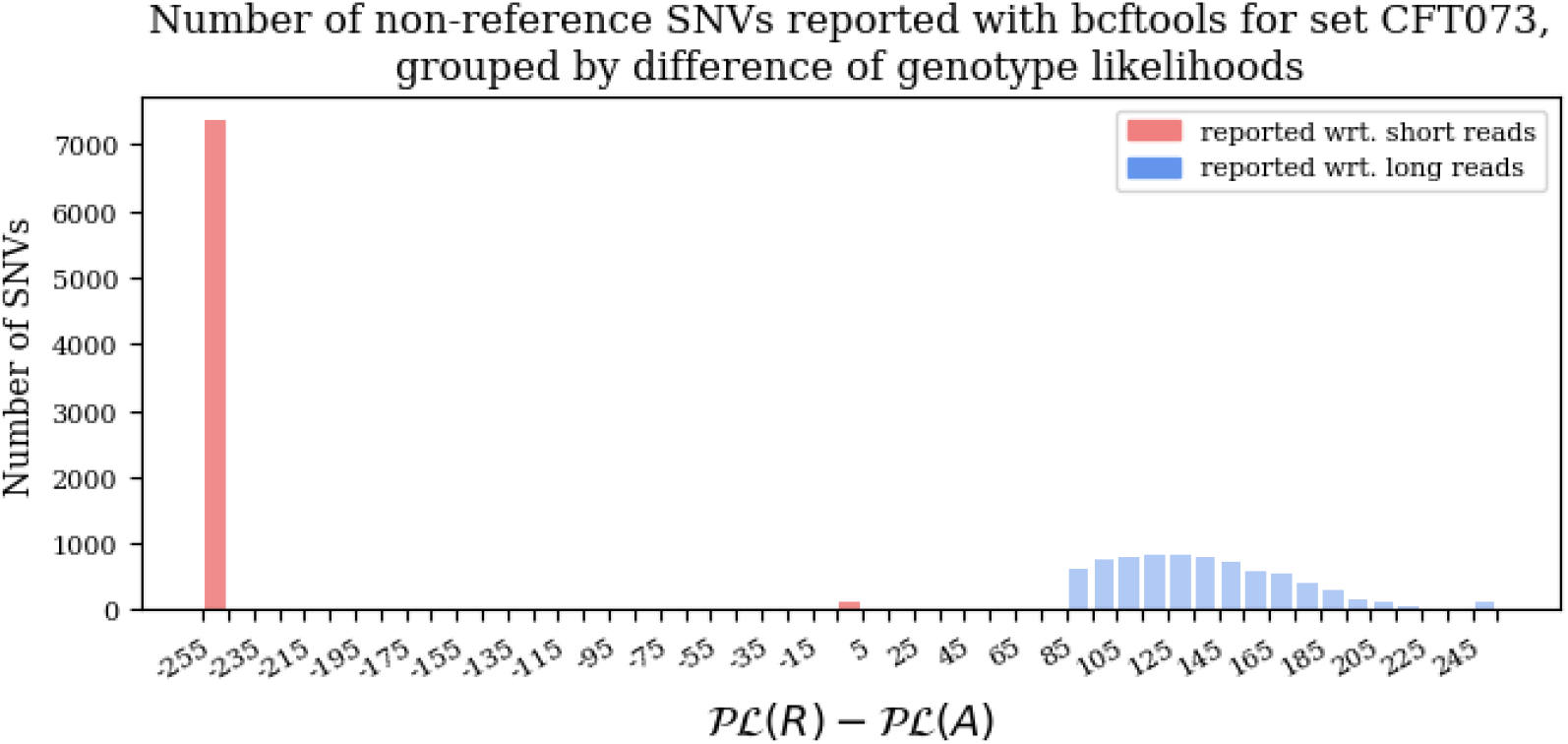
Non-reference SNVs detected with bcftools grouped by the difference of reported genotype likelihoods. The plot depicts the number of SNVs that were called with bcftools using both read sets but not contained in the set of reference SNVs. The y axis gives the number of SNVs and the x axis the difference of reported genotype likelihoods. Thereby *PL*(*R*) and *PL*(*A*) are the Phred scaled genotype likelihoods rounded to the closest integer reportet by bcftools for the reference and alternative base, respectively. A value of −255 indicates a high confidence of the reference base while a value of 255 indicates a high confidence of the alternative base.

## Discussion

In what follows the process of data selection and quality control as well as the assembly quality assessment and how the latter may have been influenced by the selected data is discussed. For each aspect limitations of this project and possible advancements for follow-up studies are pointed out.

Thereafter, the key results of this research project will be summarized and a statement on the individual assemblers and polishing tools used is given.

Based on these findings a concluding best practice recommendation is presented.

### Selection and quality control of reference data sets

A noteworthy restriction of the conducted study was the selection of real whole genome sequencing data sets exclusively. As done by other researchers, for example Wick and Holt [6] and Chen *et al*. [20], an alternative to this approach is the usage of simulated read sets. Such simulated read sets, on the one hand, allow for a highly adjustable data quality and coverage, but, on the other hand, may introduce a bias and possibly will not reflect the nature of real read data due to the underlying simulation process.

In the light of this the usage of simulated read sets seems to be superior for an accurate bench-marking but may hide issues of the assemblers being run on real data. As herein the main goal was the elaboration of a best practice applicable to real data, the exclusive usage of such data sets seems reasonable, anyway, it is suggested that the herein obtained results are validated on simulated reads. Regarding the selected data sets the quality control with FastQC but also the analysis of the BOC and per position DOC revealed several issues considered in the following.

For all long read sets, warnings regarding the read sequences quality and length distribution were raised. With respect to the fact that FastQC was especially designed for short reads, these warning can easily be explained by the underlying sequencing procedure of long reads an were therefore ignored.

The length distribution was also issued for the short read sets. This was put up with favoring a higher read coverage and the fact that only a small subset of the short reads was affected.

Warnings regarding the per position base distribution, k-mer content and GC content were given for both, long and short reads. While the GC content was validated to be normal distributed with the mean being of the respective organisms GC content, the k-mer and base distribution biases were, based on the fact that they occurred only at the starting ten read positions, thought to be due to the underlying sequencing procedure, that is to say the DNA extraction and library preparation. However, problems with the sequencing procedure or remaining adapters cannot be fully excluded. As FastQC reported no remaining adapters for the short reads and Porechop was run to trim long read adapters before starting any subsequent analysis, it seems to be more likely that the above mentioned warnings are due to the nature of sequencing process. While further trimming steps could be conducted to re-run and validate the considered assemblies, it is thought of that this will not influence the outcome of this project.

Lastly, a high duplication level of the RN4220 short reads was observed. This, however, may not be due to problems during sequencing but rather be related to the very high DOC observed. Especially as the DOC of the RN4220 reads was twice as high as for the other reference sets, but the respective reference genome is only half in length.

All of the obtained results should be considered in the light of these decisions as further data processing steps such as deduplication, quality and length filtering could affect the assemblers performance.

Concerning pre-assembly read processing, another possible step would have been the error correction of long reads. As all of the used assemblers are either especially designed for uncorrected reads or implement an internal correction step and, moreover, De Maio *et al*. were able to show that long read quality filtering can result in a decrease of the assembly quality [28] this was not conducted.

The analysis of the DOC and BOC of the read sets with respect to the selected reference genomes revealed that the CFT073 and RN4220 read sets yielded a BOC close to the expected value of 100 %. The small fraction of uncovered regions is likely to be due to true evolutionary events such as insertions and deletions. Its noteworthy that the CFT073 long reads had an BOC of 100 %, what, however, may be due to sequencing errors of the long reads as indicated by their lower quality and not vice versa. In case of the MGH78578 reference it was striking that 10 % of the reference genome was uncovered. While the major part of these uncovered regions were located on the five plasmids some segments of the bacterial chromosome of several thousands base pairs in length were revealed to have no coverage too. The fact that the major part of these uncovered segments appeared for short and long reads simultaneously strengthens the hypothesis of the originators of the reads, De Maio *et al*. [28], that these may be due to evolutionary changes of the sampled organisms over years of storage or errors of the assembly of the reference genome, especially as the latter was conducted back in 2001. These uncovered segments of the MGH78578 reference as well as the high fragmentation of the RN4220 reference genomes had a putative effect on the assemblies quality assessment:

First of all, it is apparent that no assembler would have been possible to fully reconstruct plasmids with no read coverage – this basically prevented any investigations of the capability of the assemblers to reconstruct plasmids what is suggested to be investigated on another data set.

For the MGH78578 assemblies, compared to those of the other reference sets, not only a lower genome fraction and feature recovery, but also a higher LA75 value and more misassembly events were consistently observed. This effect may be related to the nature of the reference genome as the large number of uncovered regions inevitably leads to a bias when aligning the contigs with the reference.

The assemblies of the RN4220 set yielded a high number of relocations. These were also highly conserved within all assemblies of this reference. As the reference genome is highly fragmented it is very likely that these misassembly events are not true in the sense of failures of the assemblers but rather due to aligning the contigs to the fragmented reference. These effects should definitely be taken into account when assessing the individual assemblers performances.

To tackle this presumable effect of aged or icomplete reference genomes it is suggested to choose other assessment strategies, such as a mapping of the reads back to the assemblies or a pairwise comparison of such. For the latter a variety of scoring systems and tools, like REAPR [44] or GMASS [45], could be utilized for a follow-up study.

Concerning the generated read sub samples it was successfully assessed that these yield the in theory expected DOC and BOC. Still, as the sub sampling process is a random process and especially for low mean DOCs a BOC below 100 % was achieved, this selection process may have had an influence on the assemblers performance on low coverage long reads. Especially the oscillating behaviour of some assemblers on some of the metricts, like for example the number of contigs, the LA75 value and number of misassemblies, cannot be explained unambiguously.

Thus, the generation of multiple sub samples of the same expected DOC and the assessment of the mean values for each assembler and each metric with respect to all of these could be an appropriate validation step for the findings of this project and eradicate this alleged bias of sub sampling.

Furthermore, the assessment of long read assembler performance on read sub samples was limited herein to the sub samples to the CFT073 reference set. These featured a lower read quality than the RN4220 read sets. As the assemblies of the full read sets featured a significant higher quality for the RN4220 set as for the CFT073 set, it can be assumed that the long read assemblers can even be run with read sets of lower quality than it was observed based on the results of this study. In further research one could therefore assess the long read assembler performance on down samples of the more qualitative RN4220 reads.

A further interesting consideration would be the additional sub sampling of short reads too and the assessment of the interplay of sub samples of hybrid read data.

### Assessment of assembler performance

Regarding the assessment of the assemblers performance it was basically relied on the standard procedure of investigating completeness, contiguity and correctness. As the results revealed, especially the metrics considered for completeness and contiguity did not show significant differences among assemblers for the same reference set. This, on the one hand, indicates a good performance of the assemblers regarding these aspects, but, on the other hand, might indicate an inappropriateness of the used assessment strategy.

Especially the used high coverage data already let expect almost complete assemblies and consequently one could use more stringent assessment strategies, such as the largest error free alignment or the length distribution of alignable blocks. However, these could also be determined by the nature of the reference genomes.

Furthermore, this study was not concerned with the usability of the assemblers as well as their resource requirements regarding run time and memory. This point should be considered for the best practice recommendation later in this report. At least regarding the long read assemblers these aspects are reviewed by Wick and Holt [6].

In addition, it should be noted that, if possible, default parameters were used for all the assemblers and polishing tools and that an adjustment of these parameters could not only influence the assemblers performance, be it positively or negatively, but also be the cause for the HASLR assembler either failing to complete or computing highly erroneous assemblies as well as for Trycycler being unable to be run for long read assemblies that were conducted with a coverage of below 37.58.

Concerning the assessment of the quality of assemblies that were conducted for sub sampled read sets the pattern emerged that if a large enough fraction of the reference genome was still reconstructed the correctness of the assemblies worsened. This is a logical consequence, given the fact that at this point only a hand full of reads remained per position and thus errors of these reads highly influence the assembly quality. Vice versa, if only a small fraction of the genome was reconstructed the number of errors dropped. At this point it seemed to be inappropriate to evaluate the assemblies with respect to the reference genomes any more. Under these circumstances, assemblies that dropped below a given threshold of the genome fraction should be considered as a failed assembly.

In order to enable a more appropriate evaluation of the long read assemblers performance it is suggested to run these also on sub sampled read sets with higher quality than the one of the CFT073 set. As the Unicycler hybrid assemblies were more robust towards lower long read coverages using the RN4220 set reads, this is likely to be observed for long read assemblers too.

### Applicability of long and hybrid read data for downstream applications

The applicability of long and hybrid read data for applications downstream of *de novo* genome assembly itself was assessed based on the examples of SNV discovery and genetic feature prediction.

Regarding the prediction of genetic features with GeneMarkS a clear trend showed that for low quality long reads many more and especially shorter genes were predicted than were annotated for the reference genomes. Furthermore, this effect was not observed for the assemblies of long reads with higher quality. Both observations may indicate, that sequencing errors in the long reads lead to break points of the gene prediction process, which was based on Hidden Markov Models.

The consideration that many new novel features were predicted can basically be disproved by the fact that many of the predicted genes were contained inside annotated genes, but contrary a set of non-contained genes was detected. However, this my be due to the arbitrary choice of assessing the containment as 60 % overlap and a closer investigation of the predicted genes would be necessary to fully elucidate this issue.

The discovery of SNVs was herein limited to detecting single base substitutions. The results lead to a similar conclusion as for the gene prediction: Many more SNVs were called based on long reads than based on short reads and it is unlikely that all of these are true SNVs. As a true reference SNV set was missing this hypothesis can not be proven herein. However, the fact that many SNVs had a high likelihood with respect to the long reads but an extremely low likelihood with respect to short reads as well as the higher error rate of the used long reads leads to the assumption that using long reads for SNV discovery results in a high rate of mistakenly called SNVs.

However, as the investigation of SNVs that were called using a mapping based approach had shown the long reads tended to outvote the genotype choice of the short reads and overall this may be related to a three-fold increased DOC of the long reads relative to the short reads of the DOC. To validate the results it is suggested to re-run this investigation with read sets being of equal coverage.

Furthermore it is noteworthy that only bcftools was used for this investigation and the results should be validated by using other SNV discovery methods.

Moreover, again only the CFT073 long reads with a low quality compared to the RN4220 reads were used. As the issue of feature prediction vanished for most of the RN4220 assemblies it is likely that using long reads with a high quality will also yield SNV sets with a higher agreement. This is underlined by considering that the size of the long read assembly based SNV sets basically equalled the total number of mismatches of the respective assemblies and SNV calling from long read data could greatly benefit from long reads of higher quality.

As no true reference SNV set was given and it is questionable how many of the SNVs used as reference are true SNVs or sequencing errors.

The usage of simulated data may be in addition more appropriate for this investigation and could be utilized in a more detailed study concerned with SNV discovery using long and hybrid read data.

## Conclusion

In the following, the key results of the findings of this project related to the stated research questions will be given:

Regarding general performance differences of long read and hybrid assemblers (i) it was noticeable that both classes enable the generation of highly complete and contiguous *de novo* genome assemblies. In terms of correctness a clear separation into hybrid assemblers, yielding assemblies with a very low error rate, and long read assemblers, yielding assemblies with higher error rates, was observed.

Furthermore, the long read assemblers exploiting the OLC paradigm for assembly were observed to yield lower mismatch rates in a trade off for a higher indel rate compared to *de Bruijn* based assemblers.

This separation into hybrid assemblers and more erroneous long read assemblers became remarkably less significant if long reads with a higher read quality, that is to say reads that were generated with a more recent basecaller, were used.

Given that ONT improves the accuracy of not only the sequencing devices but also the basecallers, this qualitative difference could become marginal in the near future.

The investigation of the quality of assemblies computed for sub samples of the reference read sets (ii) revealed that long read data with a depth of coverage of 60 and a mean Phred score of about 17 are sufficient to complete assemblies with, if at all, marginal differences to such assemblies computed for data featuring a higher coverage.

The hybrid assembler Unicycler was shown to be even more robust regarding low coverage long read data and enabled the computation of complete high quality assemblies with long reads featuring a mean Phred quality of 17, 20, 24 and a depth of coverage of about 40, 11 and 7 respectively.

By utilizing such low coverage or sub sampled long read sets one could possibly reduce not only the cost and run time of sequencing itself but also the computational load of the assembly tools.

Concerning the applicability of long read and hybrid assemblies or the respective read sets for applications downstream of *de novo* assembly (iii), it was revealed that the high error rate of long reads presumably influences the discovery of single nucleotide variations towards a high number of mistakenly called alternative base positions and affects the prediction of genetic features in the form of more and shorter features being predicted.

Hereafter a concluding statement on the single assemblers and polishing tools used is given.

**Canu** was able to compute complete and contiguous assemblies and featured a very low mismatch rate compared to other long read assemblers. However, the tool incorporated a high number of indel errors and tended to introduce misassemblies.

**Flye** was able to compute complete assemblies but featured a rather high mismatch rate while less indels and misassemblies were introduced compared to other long read assemblers.

**Raven** scored similar to Canu compared to the other long read assemblers but featured a less pronounced tendency to introduce misassemblies and therefore yielded more contiguous assemblies.

**Trycycler** revealed to lead to no significant improvement of the assembly quality compared to the assemblies used as input. However, by adjusting the selected assemblies a closer investigation of the tools potential could be conducted in future.

**Unicycler** was able to compute complete and contiguous assemblies on both, long and hybrid read data. While on long read data the tool did not yield significantly better results than other long read assemblers, it was able to obtain highly correct assemblies when used on hybrid read data.

**HASLR** was in this study unable to be utilized for completing any assembly without many errors. It is suggested to test the assembler with different parameters and on different data sets.

**Medaka** and **Pilon** both were revealed to lead to an improvement of the assembly correctness in the majority but not all of cases the tools were used. Especially the short read polishing had a significant effect. Possibly the polishing with the two tools can be optimized by adjusting the number of times they are run and especially Medaka, being a machine learning based tool, bears the possibility of being trained especially on data derived from the organism of interest.

### Best practice recommendation

In the light of the results obtained within this study it is recommended to run *de novo* assembly with Unicycler on hybrid read data. Thereby, a long read depth of coverage of 10 to 20 was shown to be sufficient in order to obtain highly complete, contiguous and correct assemblies, if the most recent base callers were used for read generation. The long read coverage should be increased appropriately if the quality of the long read data is lower, that is to say herein a coverage of 60 was sufficient to obtain high quality assemblies from long reads featuring a Phred quality of about 17.

As a matter of course, prior to the assembly step an appropriate quality control of the read data should be conducted.

To further tune the assembly quality it is recommended to adjust and try different parameters of the assembler, especially the number of times the internal polishing tools are run as herein it was shown that a post assembly polishing step may also decrease the assembly quality.

It remains to be investigated in future research if the coverage of short read can possibly be reduced too and how this would affect the quality of subsequent assemblies. By doing so, the computational cost of running Unicycler as well as the cost of the sequencing run may be further reduced.

## Data availability

Supplementary material and a detailed description of the methods usage are accessible via the projects GitHub repository at https://github.com/Integrative-Transcriptomics/TechnicalReport-AssemblyBestPractice.

## Author contributions statement

TH and KN conceived the idea. SH worked out the technical details and carried out the implementation. SH wrote the manuscript. All authors reviewed the manuscript.

## Acknowledgements

Funding for TH was provided by the Deutsche Forschungsgemeinschaft (DFG, German Research Foundation) Project-ID 398967434 TRR 261.

